# Plasma Protein Binding as an Optimizable Parameter for In Vivo Efficacy

**DOI:** 10.1101/2025.03.19.644079

**Authors:** Hongtao Zhao

**Affiliations:** Medicinal Chemistry, Research and Early Development, Respiratory and Immunology (R&I), BioPharmaceuticals R&D, AstraZeneca Gothenburg, Sweden

**Keywords:** pharmacokinetics, plasma protein binding, free concentrations, dose

## Abstract

Plasma protein is crucial for understanding pharmacokinetics, pharmacodynamics, and safety, yet it is often not recommended as a primary parameter for optimization in drug design. This study challenges this view by revisiting established pharmacokinetic models to elucidate a bidirectional strategy for clearance-dependent optimization of plasma protein binding. The analysis shows that strategically modulating plasma protein binding can enhance drug efficacy and safety, supporting its consideration as a key factor in the drug design process.

---

The free drug hypothesis posits that the pharmacological effect of a drug is determined by its concentration in the unbound (free) form rather than its total concentration in the body. This hypothesis is based on the understanding that only the unbound drug is able to diffuse across cell membranes, interact with target receptors, and exert a therapeutic effect. The free plasma concentration *C*_t_(free) is related to the total plasma concentration (*C*_t_) at time (*t*) by

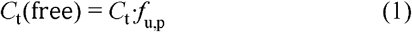

where *f*_u, p_ is the unbound fraction of the drug in plasma. This straightforward relationship might suggest that increasing the unbound fraction will enhance therapeutic effects. However, it neglects the interplay with in vivo clearance and the volume of distribution, both influencing the total plasma concentration. In other words, *C*_t_ is also dependent on *f*_u,p_. Nonetheless, the unbound area under the plasma concentration-time curve (AUC_u_) remains constant despite changes in plasma protein binding (PPB).^1, 2^ This has led to the argument that PPB should not be targeted for optimization in drug design, despite its role as a key parameter in understanding the pharmacokinetics, pharmacodynamics, and toxicities of drugs.^1, 2^ Here, we unveil its opposing optimization directions in the context of high and low intrinsic clearance, adjusted for the volume of distribution.

For a drug administered as a single intravenous dose following first-order elimination in a one-compartment model, *C*_t_ is given by

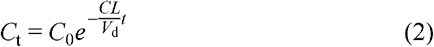

where *C*_0_ is the initial plasma concentration, *CL* is the in vivo clearance and *V*_d_ is the volume of distribution. *C*_0_ is determined by

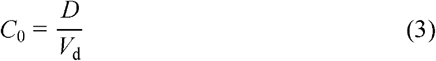

where *D* is the administered dose. According to the well-stirred model,^3^ in vivo clearance (*CL*) is influenced by liver blood flow (*Q*), the unbound intrinsic clearance (*CL*_int,u_), and *f*_u,p_

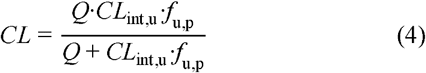

Assuming *Q* >> *CL*_int, u_· *f*_u, p_,

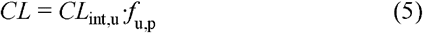

The volume of distribution (*V*_d_) reflects the relative binding affinity of a drug between plasma and tissues^4^

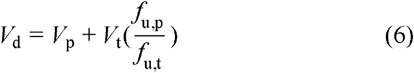

where *V*_p_ is the plasma volume, *V*_t_ is the apparent tissue volume, and *f*_u,t_ is the unbound fraction in tissue. The unbound area under the plasma concentration-time curve (AUC_u_) can be determined by integrating the unbound plasma concentration over time

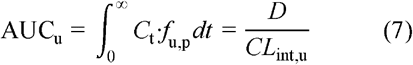

As such, AUC_u_ remains constant despite changes in plasma protein binding, meaning that the average free plasma concentration is unaffected.^1^ However, this does not mean that time-dependent free concentrations (e.g., free *C*_max_) remain unchanged, which is a misconception that could occur.^5^

To assess the effect of the unbound fraction in plasma (*f*_u,p_) on the free plasma concentrations, it is essential to examine its correlation with the volume of distribution (*V*_d_). An analysis of 4,167 rat pharmacokinetic data points against rat plasma binding measurements at AstraZeneca reveals no correlation, suggesting a strong correlation between *f*_u,p_ and *f*_u,t_ (**Figure 1**). The lack of clear correlation has been demonstrated by nine analogues of 5-*n*-alkyl-5-ethyl barbiturate derivatives, where changing the alkyl group from methyl to nonyl resulted in *f*_u,p_ changes from 0.01 to 1, while *V*_d_ remained largely unchanged.^6^ Assuming a constant *V*_d_, time-dependent free plasma concentrations were simulated at two different unbound fractions, 1% and 10%, starting with an initial total plasma concentration of 1 μM. Additionally, both high and low intrinsic clearance, adjusted for the volume of distribution (*CL*_int,u_/*V*_d_ = 5 and 0.5 h^-1^, respectively), were considered (**Figure 2**).

**Figure 1.**
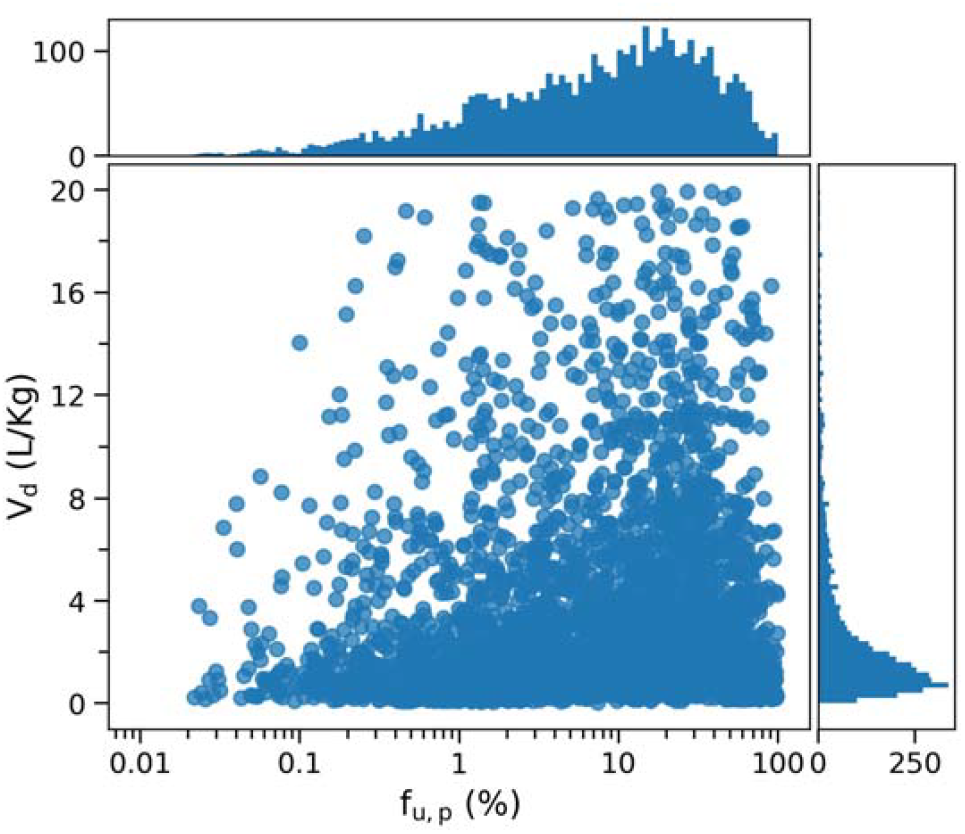
Scatter plot showing the volume of distribution (*V*_d_) at steady state against the unbound fraction in plasma (*f*_u,p_), measured in male Han Wistar rats.

**Figure 2.**
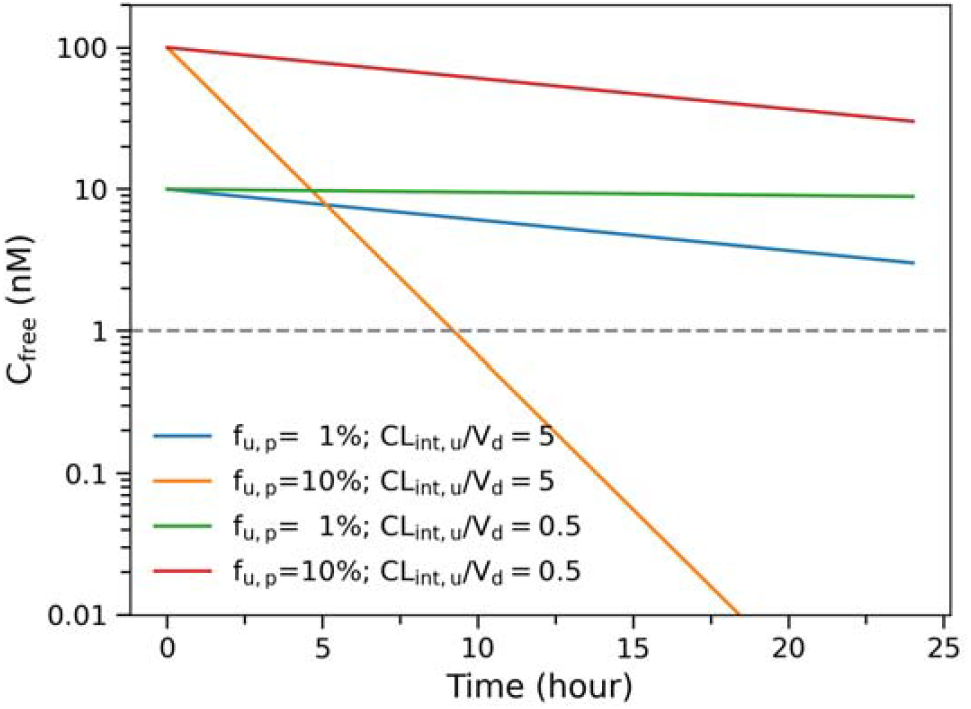
Simulated free plasma concentration-time curves at an initial total plasma concentration of 1 μM assuming a constant *V*_d_.

At a high clearance (*CL*_int,u_/*V*_d_ = 5 h^-1^), the free plasma concentration at an unbound fraction of 1% decreases from 10 nM to 3 nM over 24 hours, with a half-life of 13.9 hours. Conversely, at a 10% unbound fraction, the concentration decreases from 100 nM to 0.6 pM in the same period, with a half-life of 1.4 hours. Assuming a minimum therapeutic concentration of 1 nM, more frequent dosing would be necessary. Thus, for high *V*_d_-adjusted clearance, increasing plasma protein binding could be a viable strategy to optimize the effective half-life of drug candidates in humans. This strategy is exemplified by optimizing acidic CXC chemokine receptor 2 (CXCR2) antagonists, which have a low *V*_d_ of 0.3 L/kg.^7^

At a low clearance (*CL*_int,u_/*V*_d_ = 0.5 h^-1^), the free plasma concentration at an unbound fraction of 1% decreases from 10 nM to 9 nM after 24 hours, with a half-life of 139 hours. In contrast, at a high unbound fraction of 10%, the free plasma concentration decreases from 100 nM to 30 nM after 24 hours, with a half-life of 13.9 hours. In such cases, reducing plasma protein binding can mitigate the risk of drug accumulation and associated toxicity.

The effect of plasma protein binding on pharmacokinetics could be better illustrated from a mathematical perspective upon explicitly rewriting **Eq. 1** as

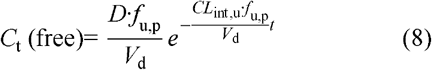

Intrinsic clearance appears only in the exponent, making its optimization straightforward; reducing it is always beneficial for increasing free drug concentrations.^2^ In contrast, the volume of distribution (*V*_d_) and the unbound fraction in plasma (*f*_u,p_) appear both in the exponent and the pre-exponential term. As such, changes in *V*_d_ or *f*_u,p_ do not impact the average free plasma concentration but rather influence the effective half-life, which determines the dosing regimen. Since the average free plasma concentration remains constant, the total amount of drug administered during treatment can remain the same by adjusting dosage and frequency. Optimization of *V*_d_ or *f*_u,p_ for an effective half-life is bidirectional, depending on intrinsic clearance. Given that *V*_d_ and *f*_u,p_ are reciprocal, the unbound volume of distribution (*V*_d,u_ = *V*_d_/*f*_u,p_) is more insightful than either parameter individually.^5^ Unlike *V*_d_, *f*_u,p_ can be routinely measured, and due to the lack of correlation between them, *f*_u,p_ could be a more practical parameter for optimizing half-life. In small molecule drug discovery, lead compounds frequently have medium to high intrinsic clearance, making it challenging to further reduce it. Thus, increasing plasma protein binding or *V*_d_ provides an alternative approach for achieving an effective half-life.

In summary, plasma protein binding is a crucial parameter for optimizing the effective half-life of drugs in the discovery phase. Its optimization should be tailored according to the intrinsic clearance of the compound. For drugs with low intrinsic clearance, reducing plasma protein binding is beneficial to prevent drug accumulation and potential toxicity. Conversely, for those with high intrinsic clearance, increasing plasma protein binding can help achieve a desired effective half-life. This approach ensures that the pharmacokinetic profile is optimized to enhance the therapeutic potential of drug candidates.

## Notes

H.Z. is an employee of AstraZeneca and may own stock or stock options.

## TOC

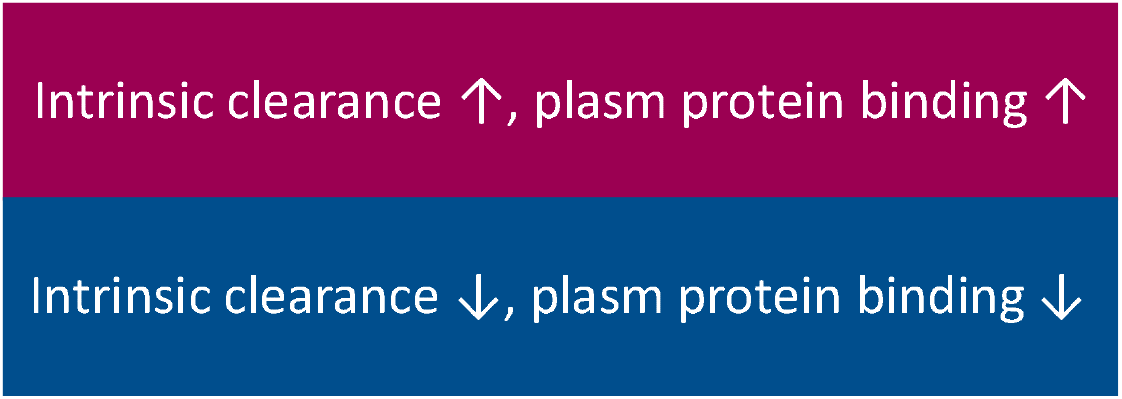

